# Naturalistic Movie Viewing Reveals a Distributed Network Supporting Audiovisual Speech Processing

**DOI:** 10.64898/2026.10.26.762007

**Authors:** Hailey C. Smith, Ryan M. O’Leary, Jonathan E. Peelle

**Affiliations:** Institute for Cognitive and Brain Health, Northeastern University, Boston, MA 02115; Department of Psychology, Northeastern University, Boston, MA 02115; Department of Communication Sciences and Disorders, Northeastern University, Boston, MA 02115

**Keywords:** Neuroimaging, Audiovisual Integration, Speech, fMRI

## Abstract

Traditional paradigms in cognitive neuroscience often fail to capture the richness of real-world experiences. To bridge this gap, researchers increasingly use naturalistic stimuli such as movies, which provide complex, multisensory input. Speech perception is a prime example of such complexity: it is inherently multisensory, involving the integration of auditory cues, lip movements, and facial expressions. Prior studies suggest that the left posterior superior temporal sulcus (pSTS) and superior temporal gyrus (STG) play a critical role in this integration. However, most studies of audiovisual speech have relied on simplified or artificial stimuli, limiting ecological validity. To address these gaps, we analyzed fMRI data from human participants watching two full-length films (*500 Days of Summer, Citizenfour*) to assess the consistency of multisensory speech processes across distinct genres and narratives. We manually annotated word onset times and modality (audiovisual vs. auditory-only). Across two movies and 38 participants we observed consistent speech-related activation in the pSTS and STG across speech types. Compared to auditory-only speech, audiovisual speech recruited a broad left-lateralized network including the angular gyrus, posterior temporal cortex, temporal pole, and inferior frontal gyrus, along with the right cerebellum. These findings were also consistently present in single-subject data. Importantly, most of the regions observed in our analyses have been implicated in multisensory function in the nonhuman primate literature despite receiving relatively little focus in human neuroimaging. Together our results demonstrate that speech processing during naturalistic viewing recruits a distributed network extending substantially beyond regions traditionally emphasized in audiovisual speech research.

## Introduction

Real-world spoken communication is rapid, contextually constrained, and shaped by environmental variability. These elements create opportunities to investigate how our senses collaborate in everyday environments, making speech perception an ideal vehicle for studying multisensory integration. Audiovisual speech exemplifies this process, combining auditory signals with visual information from the speaker, such as lip movements, facial expressions, and body gestures. The availability of complementary auditory and visual cues allows listeners to extract meaning more efficiently from audiovisual than from unimodal speech, especially in challenging listening environments (Sumby & Pollack, 1954). Consequently, understanding how the brain combines auditory and visual speech signals is crucial for explaining how people successfully navigate complex, real-world conversations.

Research in non-human primates indicates that regions within the superior temporal sulcus (STS) respond to information from multiple sensory modalities including auditory and visual input (Benevento et al., 1977; Bruce et al., 1981). Subsequent work has demonstrated that multisensory communication processing extends across a broader network of temporal and prefrontal regions, including the anterior temporal lobes and ventrolateral prefrontal cortex (Khandhadia et al., 2021; Romanski, 2007). In humans, functional neuroimaging studies have identified the posterior superior temporal sulcus (pSTS) as a key hub for combining auditory and visual speech signals (Beauchamp et al., 2004; Beauchamp et al., 2002; Poremba et al., 2003; Stevenson et al., 2011). Classic experimental paradigms, such as the McGurk illusion (McGurk & MacDonald, 1976), demonstrate that conflicting auditory and visual speech cues alter perception and engage the pSTS, providing evidence for its role in multisensory binding (Beauchamp et al., 2004, 2010). In non-McGurk stimuli, in which auditory and visual cues are always congruent, activity increases in several speech-sensitive regions, including the superior temporal gyrus (STG) and other visual processing areas (Calvert et al., 2000, 1999; Peelle, 2022).

Traditional paradigms have substantially advanced our understanding of the neural mechanisms supporting speech processing, often using controlled auditory and audiovisual stimuli such as isolated phonemes, words, simple sentences, and visual speech cues to elicit responses in the STG and related auditory networks. However, elucidating the neural basis of speech processing in real-world settings may require paradigms that better capture the complexity of everyday communication generally (Shamay-Tsoory and Mendelsohn, 2019; Sonkusare et al., 2019), extending to audiovisual speech perception (Van Engen et al., 2022). In particular, the degree to which multisensory integration mechanisms identified in controlled paradigms generalize to naturalistic communication is unclear.

To address this gap, many researchers now adopt naturalistic paradigms with continuous, dynamic, and context-rich stimuli to better reflect brain processing in real life (Huth et al., 2016; Chen et al., 2017; Aliko, 2020; Sherafati et al., 2025). These methods, specifically movie-watching, elicit reliable sensory responses across people, especially in visual, auditory, and higher-order regions (e.g., Hasson et al., 2004). Because speech in real-world environments occurs within rich visual and narrative contexts, movie watching provides an opportunity to examine how auditory and visual speech cues are processed during continuous communication. Further, having these rich stimuli also enables investigation of audiovisual variability in dynamic environments, as movies naturally include a combination of visual speakers, off-screen speech, and background noise, elements often missing from traditional experiments. Despite the growing use of naturalistic stimuli to understand speech comprehension, relatively little work has tested neural systems supporting audiovisual speech perception in naturalistic contexts.

Our study addresses this gap by using functional magnetic resonance imaging (fMRI) data from participants watching one of two full-length films from the Naturalistic Neuroimaging Database (NNDb; Aliko et al., 2020). For each movie, we manually annotated spoken words in the films as audiovisual (speech input with corresponding face and mouth movement on screen) or auditory-only (speech input without corresponding face and mouth movement on screen).

Based on prior literature, we predicted that canonical auditory and audiovisual speech regions (e.g., STG, pSTS, visual cortex) would differentially respond to auditory-only and audiovisual speech. Importantly, we also hypothesized that this functional organization would remain robust under naturalistic viewing conditions, reflecting generalizable multisensory integration mechanisms that extend beyond simplified experimental paradigms.

## Method

### Participants

Participants consisted of a total of 38 adults (19 females, age range of 19-58 years, M=27.36, SD=10.18) from the NNDb (Aliko et al., 2020). All participants reported being native English speakers, right-handed, without hearing impairment, with normal or corrected-to-normal vision, and no history of neurological or psychiatric illness. All participants provided informed consent in line with the institutional review board guidelines. More detailed demographic information can be found in Aliko et al. (2020).

### Movie Procedure

Participants viewed either *500 Days of Summer* (2009; duration 91.17 minutes; n=20) or *Citizenfour* (2014; duration 113.4 minutes; n=18) during continuous fMRI acquisition, corresponding to approximately 64 hours and 14 minutes of fMRI data across participants. The movie was presented using a rear-projected screen and viewed through a mirror mounted on the head coil. Audio was connected through MR-compatible headphones and each film was presented continuously, with participants permitted to take breaks as needed. During breaks, movie playback was paused and subsequently resumed in synchronization with the scanner trigger pulses. Participants were asked to remain as still as possible and to watch the movie as they normally would.

### Speech Annotation

The NNDb provides transcripts and approximate word timings generated through automated speech recognition and manual correction. However, because automated transcription can contain errors and does not distinguish between audiovisual and auditory-only speech, we manually verified the transcripts, corrected inaccuracies, and annotated the timing of each word onset in the films. Audiovisual speech events were defined as words for which visual speech information from the speaker was clearly available. This required (1) visible articulation synchronized with the spoken word, (2) an unobstructed view of the speaker’s mouth, (3) a camera angle providing a clear view of the face, and (4) at least 50% of the speaker’s face visible on screen. Words were classified as auditory-only when these criteria were not met and the mouth of the speaker producing the word was either off-screen or not visible. All words and onset events were then cross-checked between raters. The annotation pipeline is illustrated in Figure 1. Across the two films, we annotated 8,378 speech events in *500 Days of Summer* (5,007 audiovisual, 3,371 auditory-only), and 14,258 events in *Citizenfour* (7,549 audiovisual, 6,709 auditory-only).

**Figure 1:**
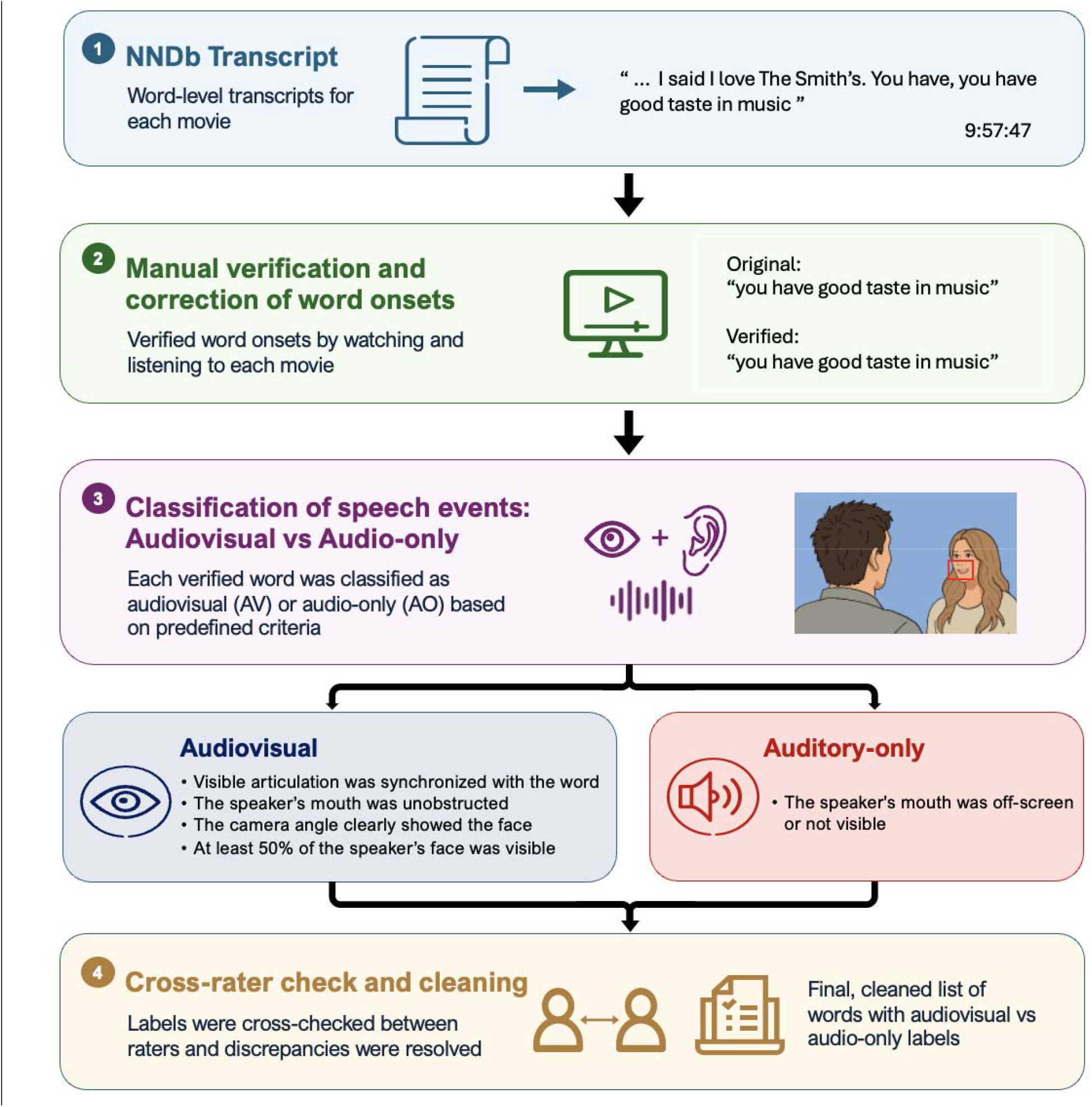
Audiovisual word classification. Overview of word onset verification and classification of speech events as audiovisual (AV) or auditory-only (AO)

### fMRI Acquisition and Preprocessing

Full details on data acquisition and preprocessing are given in Aliko et al. (2020). In brief, data were acquired using a 1.5 T Siemens MAGNETOM Avanto scanner using a 32-channel head coil. Functional images were obtained using a multiband echoplanar imaging (EPI) sequence (TR = 1 s, TE = 54.8 ms, flip angle = 75 degrees, 40 interleaved slices, 3.2 mm voxel size). T1-weighted anatomical images were acquired using a 10-minute MPRAGE sequence (TR = 2.73 s, TE = 3.57 ms, 1 mm resolution). Images were preprocessed using AFNI and FSL tools; functional data were corrected for slice-timing differences, despiked, and motion-corrected by aligning each volume to a reference volume. Functional images were coregistered to each participant’s anatomical image and normalized to MNI space. Spatial smoothing was applied to a target of 6 mm. Time series were detrended and cleaned using nuisance regression, including motion parameters, white matter signal, cerebrospinal fluid signal, and low-frequency drift. Independent component analysis was used to identify additional artifacts that were regressed out of the data.

### fMRI Modeling

At the single-subject level, neural responses were modeled using a general linear model (GLM) implemented in SPM12 revision 7770 (Wellcome Centre for Human Neuroimaging; RRID:SCR_007037). Word onset times obtained from the speech annotation procedure were used to construct event regressors corresponding to audiovisual and auditory-only speech events. Each event was modeled as a stick function at the time of word onset and convolved with SPM’s canonical hemodynamic response function (HRF) to generate predicted responses. Additional regressors were included to account for visual and stimulus-related confounds (see Control Regressors section). First-level GLMs were estimated separately for each subject using the same set of stimulus-locked regressors. From the fitted GLM, first level contrast images were generated to assess neural responses to audiovisual and auditory-only speech. Primary contrasts included auditory-only, audiovisual, and audiovisual > auditory-only.

#### Control Regressors

While not the primary focus of the present study, we wished to ensure that observed differences in brain activity between auditory-only and audiovisual speech were not the result of low-level confounds. For this purpose, we included a suite of regressors to account for visual changes using functions from the OpenCV computer vision model (Bradski, 2000). These included measures of overall luminance and contrast, color properties in HSV space, edge and texture statistics (Sobel edge magnitude and Laplacian variance), optical-flow estimates of motion energy, and shot boundary detection. First temporal derivatives of luminance and contrast were included to index rapid visual changes. Additional regressors were included for higher-level visual confounds such as camera motion, as well as the presence and number of faces. These values were computed for each frame in the movie, convolved with SPM’s canonical hemodynamic response function (HRF), and downsampled to 1-second TRs to create regressors that were temporally aligned to the BOLD signal. Shot boundaries were treated as impulse-like events at TRs, and were also convolved before entering the model. We also included the auditory amplitude envelope as an additional regressor.

#### Group-level Analysis

Group-level analyses were conducted using nonparametric permutation testing implemented in FSL randomise (Smith et al., 2004). Individual subject contrast images were entered into a one-sample permutation test using 5000 permutations with threshold-free cluster enhancement (TFCE) to correct for multiple comparisons across voxels (Smith & Nichols, 2009). A group mask was created to restrict analyses to voxels that were present in all of the subjects. The resulting statistical maps were thresholded at p < .05 (FWE-corrected).

To quantify the spatial consistency of activation across the two movies, Dice similarity coefficients were calculated separately for each contrast. Thresholded statistical maps (TFCE, FWE-corrected, p < .05) were converted to binary masks, and the number of voxels surviving the statistical threshold in each movie-specific map, as well as the number of overlapping voxels, were obtained using FSL via fslstats. Dice similarity coefficients were then calculated as:

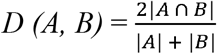

where *A* and *B* represent the thresholded voxel masks for *500 Days of Summer* and *Citizenfour*, respectively.

## Results

We first examined neural responses to auditory-only (AO) and audiovisual (AV) speech relative to baseline to determine whether spoken words elicited activation within canonical speech-processing regions. As shown in Figure 2, both AO and AV speech elicited robust bilateral activation in the superior temporal gyri (STG). Activation extended into the posterior superior temporal sulcus (pSTS), inferior frontal gyrus (IFG), superior frontal gyrus (SFG), supplemental motor area (SMA) and right cerebellar regions including Crus I and II. More extensive pSTS activation was observed during AV speech. Overlap maps demonstrated consistent activation patterns across *500 Days of Summer* and *Citizenfour* for both AO (Dice coefficient = 0.69) and AV (Dice coefficient = 0.65) speech, indicating that the overall spatial distribution of speech-related activation was similar across the two naturalistic movies.

**Figure 2.**
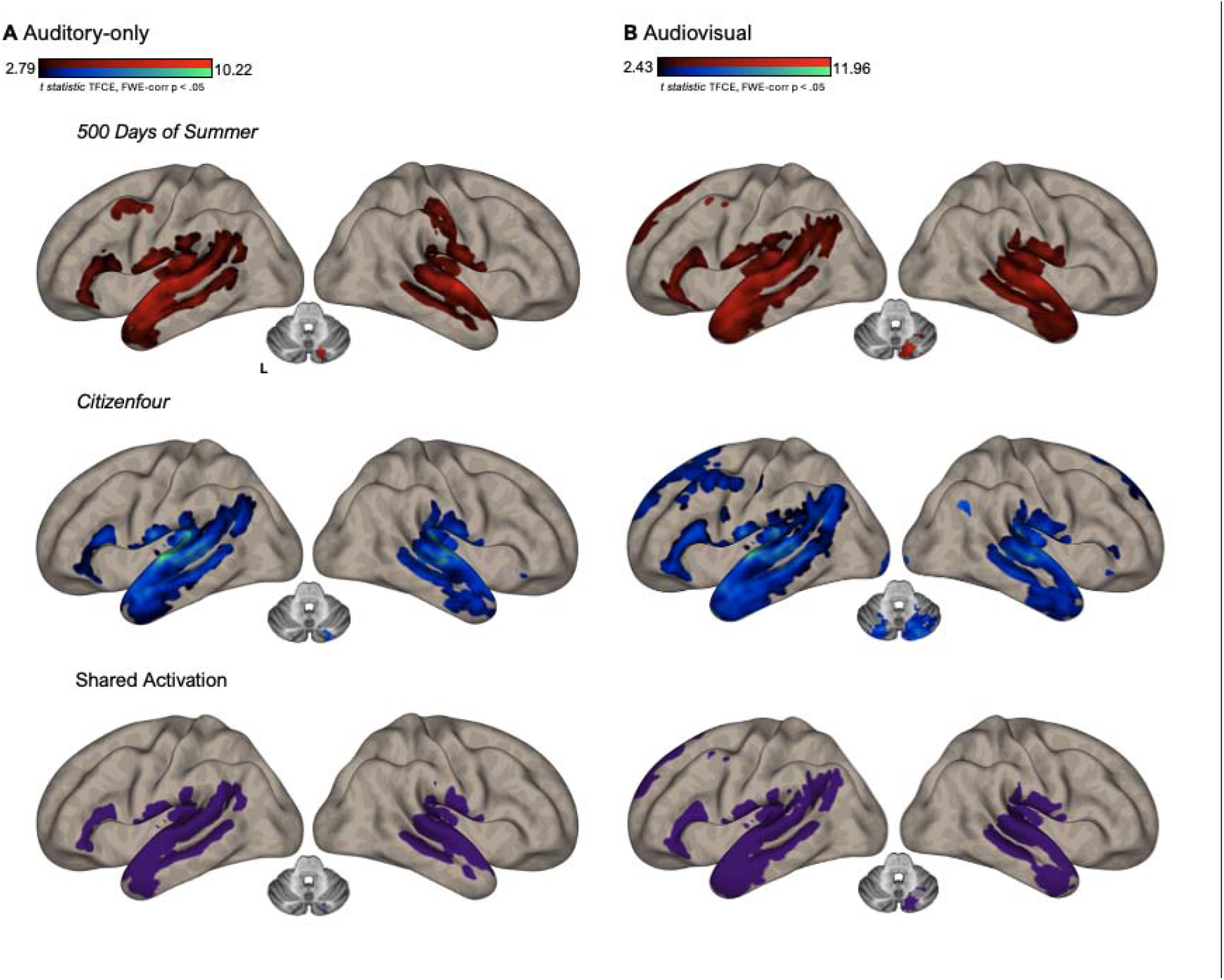
Baseline speech responses during naturalistic viewing for auditory-only (A) and audiovisual (B) words in *500 Days of Summer* (red) and *Citizenfour* (blue). Purple binary overlap maps show activation shared across the two movies. Brain maps are displayed on left and right lateral inflated surfaces. Axial slices show cerebellar activation and are displayed at MNI z = -31. Statistical maps were thresholded using threshold-free cluster enhancement (TFCE) with family-wise error (FWE) correction at p < .05.

We next examined regions that showed greater activation during audiovisual (AV) speech than auditory-only (AO) speech. In *500 Days of Summer*, the AV > AO contrast revealed bilateral activation encompassing superior and middle temporal gyri, anterior temporal pole, inferior frontal cortex, angular gyrus, superior/medial frontal cortex, and supplementary motor regions. In *Citizenfour*, the response was more strongly left-lateralized and involved middle and anterior temporal cortex, left inferior frontal cortex, left angular gyrus, superior/medial frontal cortex, and supplementary motor area. There was additional activation in middle frontal, occipital, inferior temporal, postcentral, and cerebellar regions. Cerebellar activity included Crus I and II, lobules III, IV, VIIb, and VIII, and vermal lobules IV–V, VIII, and IX. Cross-movie overlap was concentrated primarily in a left-lateralized network involving middle/anterior temporal, inferior frontal, angular, and medial frontal/supplementary motor regions (Dice coefficient = 0.26) as seen in Figure 3. Notably, no significant visual activation was observed for audiovisual speech, either relative to baseline or relative to auditory-only speech.

**Figure 3.**
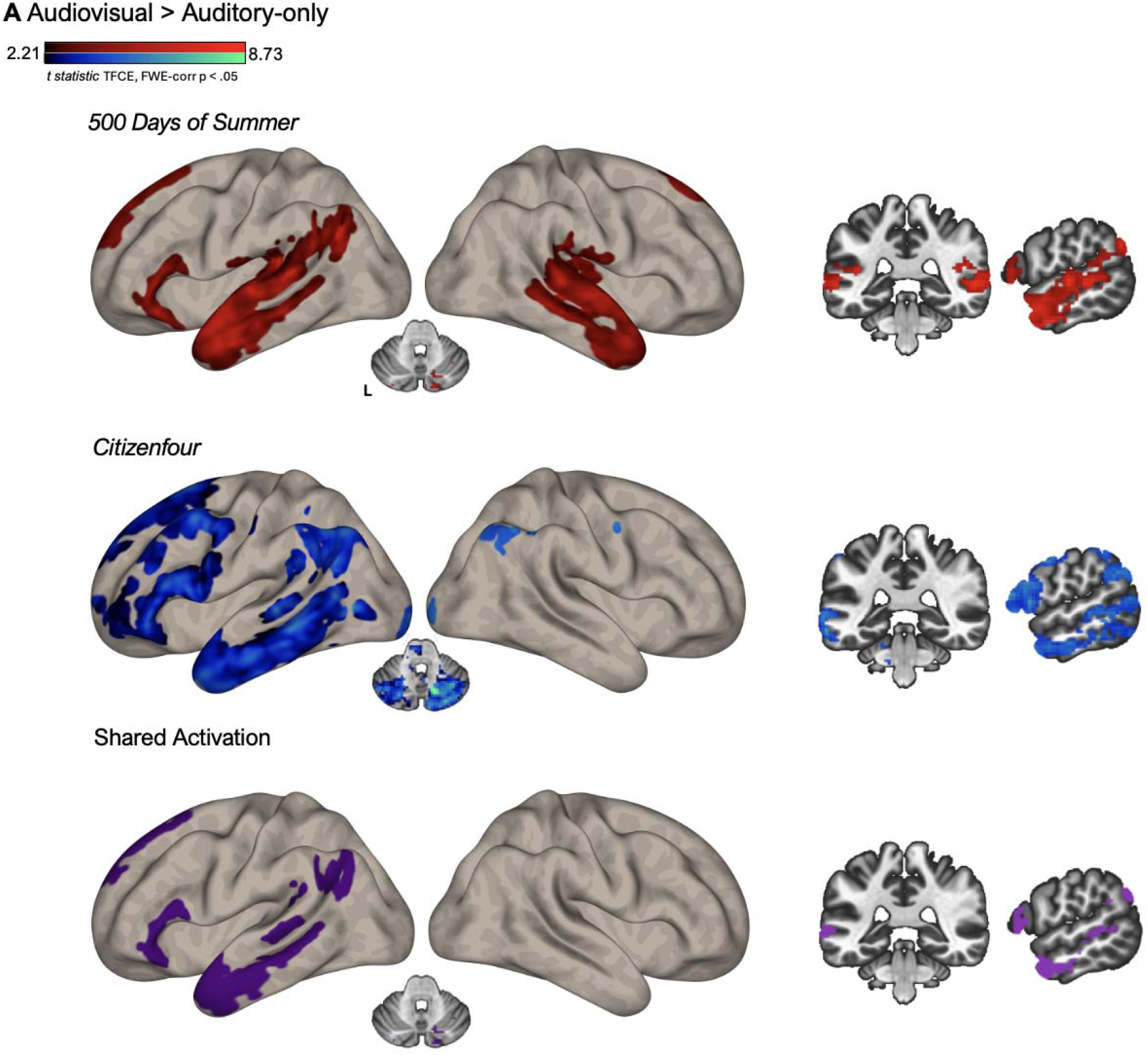
Audiovisual > auditory-only (AV > AO) contrast maps during naturalistic movie viewing for *500 Days of Summer* (red) and *Citizenfour* (blue). Purple binary overlap maps show activation across the two movies. Cortical maps are displayed on the left and right lateral inflated surfaces. Coronal (MNI y = -36) and sagittal (MNI x = -57) slices provide additional views of activation not fully represented on the lateral cortical surfaces. Axial slices show cerebellar activation and are displayed at MNI z = -31. Statistical maps were thresholded using threshold-free cluster enhancement (TFCE) with family-wise error (FWE) correction at p < .05.

To ensure that the group-level results were also evident in individual participants, we examined single-subject activation maps from all 38 participants to determine whether activity was seen in each of the primary regions (Devlin & Poldrack, 2007). A representative group of participants is shown in Figure 4. Activity was observed in the pSTS, IFG, and medial frontal regions in every participant (38/38), as well as the angular gyrus (37/38) and the temporal pole (35/38). Although the spatial extent of activation varied across individuals, the principal regions identified at the group level were evident in the overwhelming majority of individual participants.

**Figure 4.**
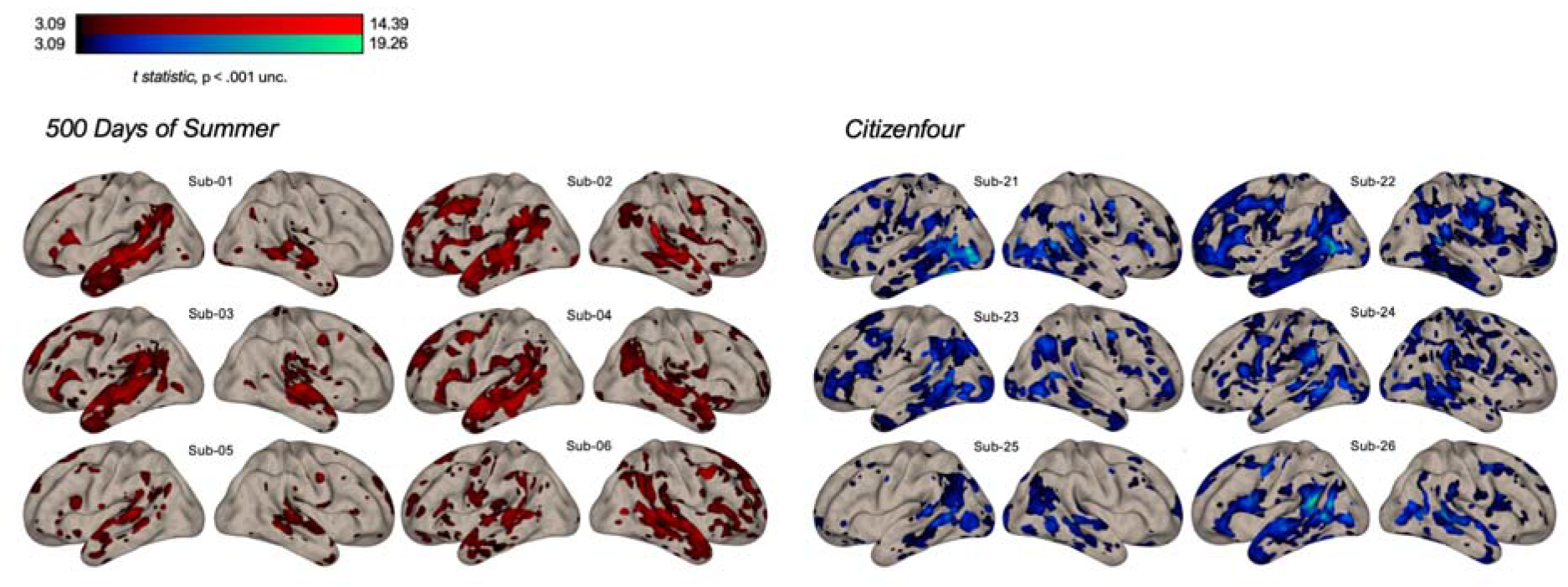
Example single-subject audiovisual > auditory-only (AV > AO) contrast responses during naturalistic viewing for *500 Days of Summer* (left; red) and *Citizenfour* (right; blue). Cortical maps are displayed on left and right lateral inflated surfaces at p < .001 uncorrected. Subject labels correspond to NNDb subject IDs.

## Discussion

In the present study we investigated speech processing when communication is embedded in a realistic environment. We found that canonical speech-processing regions were robustly recruited during both auditory-only and audiovisual speech across two distinct movies. Although the additional neural recruitment associated with audiovisual speech varied between movies, a common left-lateralized network spanning middle and anterior temporal, inferior frontal, and medial frontal regions was observed across both *500 Days of Summer* and *Citizenfour*.

Our findings are consistent with previous work demonstrating that left STG and pSTS play central roles in audiovisual speech perception and multisensory integration (Bruce et al., 1981; Peelle et al., 2022; Beauchamp et al., 2004). In the present study, baseline audiovisual speech robustly recruited posterior temporal regions, including STG and pSTS, across two independent movies. Although many previous demonstrations of audiovisual integration have relied on controlled paradigms, such as the McGurk illusion or single words, naturalistic neuroimaging studies suggest that canonical language and speech systems remain engaged when speech is embedded within continuous, context-rich stimulation (Jääskeläinen et al., 2021). Contrary to our initial prediction, compared to auditory-only speech, audiovisual speech did not elicit significant additional recruitment of the visual cortex. This may reflect an important distinction between controlled audiovisual speech paradigms and naturalistic movie viewing. In controlled paradigms, conditions containing visual speech information are often compared with conditions containing little or no meaningful visual information, whereas in the present study, continuous visual input was present throughout both auditory-only and audiovisual speech. Thus, the presence of visible articulatory information during audiovisual speech may not produce additional visual cortical recruitment beyond that already elicited by the ongoing visual environment. It is also important to consider the presence of acoustic noise from the scanner, which may influence multisensory processing (Peelle, 2014).

In contrast to the consistency of baseline speech responses across movies, regions preferentially recruited during audiovisual speech exhibited lower spatial overlap. Naturalistic stimuli contain multiple, dynamically changing sensory and contextual features and can therefore engage several functional systems simultaneously (Saarimäki, 2021). One possible explanation for this increased variability is that audiovisual speech engages additional frontal and sensorimotor systems whose recruitment is more sensitive to the contextual features that vary across movies, such as conversational context, vocabulary, and emotional engagement. Although the precise spatial extent of AV > AO recruitment differed between narratives, cross-movie overlap revealed a common left-lateralized network extending across temporal, frontal, and parietal regions (Figure 3).

Importantly, the audiovisual-specific recruitment we observed during movie viewing extended beyond the posterior temporal regions traditionally emphasized in human audiovisual speech research. These observations are consistent with data from neurophysiological studies in non-human primates indicating that multisensory communication processing extends beyond pSTS. For example, single-unit recordings during face-voice communication show audiovisual interactions not only in STS but also in anterior voice-sensitive temporal regions (Perrodin et al., 2014). Further work examining the face patch system similarly suggests interactions between anterior temporal face processing and auditory communication signals (Khandhadia et al., 2021). The temporal pole itself has also been proposed as a higher-order convergence region in primates, with macaque temporal-pole subdivisions receiving input from both auditory and visual temporal association cortex and showing broad comparative correspondence with human temporal-pole regions (Insausti, 2013).

Additional work in non-human primates has identified multisensory responses to faces and vocalizations in the ventrolateral prefrontal cortex (VLPFC) that researchers explicitly relate to the human inferior frontal cortex. The VLPFC contributes to the processing and maintenance of those communication signals (Romanski, 2007; Romanski, 2012; Plakke, 2015). Importantly, these distributed multisensory responses are not limited to static or highly simplified stimuli, with similar temporal and prefrontal involvement reported during more naturalistic audiovisual stimulation in non-human primates (Diehl & Romanski, 2014).

The angular gyrus provides another potential point of correspondence with non-human primate multisensory networks. Comparative anatomical work identifies macaque area PG as a likely homologue of human PGa, with macaque Opt similarly corresponding to human PGp (Niu & Palomero-Gallagher, 2023). Importantly, area PG receives convergent input from visual, auditory, and somatosensory association regions, supporting a broader role for posterior inferior parietal cortex in multisensory processing (Rozzi et al., 2006). The medial frontal recruitment observed here may also reflect engagement of sensorimotor systems that link incoming communication signals to action. In non-human primates, SMA neurons encode auditory information during sound categorization and transition from acoustic to movement-related representations, suggesting a role in transforming auditory signals into behaviorally relevant motor plans (Morán et al., 2021). SMA activity has also been observed preceding vocalization, supporting a role in vocal preparation and control (Gemba et al., 1997). Related work in preSMA suggests a broader role in motor preparation during volitional vocal behavior rather than call-specific initiation or timing (Gavrilov et al., 2017).

Together, recruitment of anterior temporal, inferior frontal, and parietal regions in the present study may reflect engagement of a broad multisensory communication network under naturalistic viewing conditions. The specific role of these regions in audiovisual speech processing will require additional investigation.

Several limitations should be considered when interpreting the present findings. First, although the use of two feature-length films allowed us to examine cross-movie consistency, the small number of movies limits generalization across the broader range of naturalistic narratives. Future work incorporating a larger and more diverse set of films could determine which responses to speech remain consistent across genres, speakers, and visual contexts. Audiovisual speech labels reflected the availability of visible articulatory information on screen rather than whether participants actually fixated the speaker’s mouth. Relatedly, the binary classification of words as auditory-only or audiovisual necessarily simplifies the continuous variability in visual speech cues that occurs during natural viewing. Finally, although movies provide a more naturalistic stimulus than many conventional laboratory paradigms, fMRI acquisition still constrains ecological validity through scanner noise, restricted movement, and passive viewing.

Future studies could address these limitations by combining naturalistic speech paradigms with eye tracking and more detailed models of visual speech availability, as well as by sampling a broader range of films and speakers. Quieter or more portable neuroimaging methods, including fNIRS or EEG, may also enable speech processing to be examined under less constrained listening conditions (Von Lühmann, 2021). Ultimately, extending naturalistic speech research from passive movie viewing to live conversation may provide the clearest test of how these distributed networks operate during everyday communication, when listeners must not only comprehend speech but also anticipate, respond, and coordinate with another person.

In conclusion, the present study demonstrates that canonical speech-processing networks remain robust across distinct naturalistic narratives. Although the temporal organization of auditory and audiovisual speech was consistent between movies, additional recruitment associated with audiovisual speech varied across contexts. Nevertheless, shared audiovisual-specific responses extended into anterior temporal and frontal regions, consistent with a distributed network supporting multisensory communication. These findings highlight both the stability and flexibility of the neural systems supporting multisensory speech perception and demonstrate the value of naturalistic paradigms for examining how these systems operate within complex, context-rich communication.

## Acknowledgments

Work reported here was supported in part by grant R01 DC019507 from the US National Institutes of Health. We are grateful to the undergraduate research assistants in our lab for their extensive contributions to the annotation and word-labeling efforts that supported this work.

## References

Aliko, S., Huang, J., Gheorghiu, F., Meliss, S., & Skipper, J. I. (2020). A naturalistic neuroimaging database for understanding the brain using ecological stimuli. Scientific Data, 7(1), 1–21. 10.1038/s41597-020-00680-2

Ashburner, J., Barnes, G., Chen, C. C., Daunizeau, J., Flandin, G., Friston, K., … & Penny, W. (2014). SPM12 manual. Wellcome Trust Centre for Neuroimaging, London, UK, 2464(4), 53.

Beauchamp, M. S., Lee, K. E., Haxby, J. V., & Martin, A. (2002). Parallel Visual Motion Processing Streams for Manipulable Objects and Human Movements. Neuron, 34(1), 149–159. 10.1016/S0896-6273(02)00642-6

Beauchamp, M. S., Argall, B. D., Bodurka, J., Duyn, J. H., & Martin, A. (2004). Unraveling multisensory integration: Patchy organization within human STS multisensory cortex. Nature Neuroscience, 7(11), 1190–1192. 10.1038/nn1333

Beauchamp, M. S., Lee, K. E., Argall, B. D., & Martin, A. (2004). Integration of Auditory and Visual Information about Objects in Superior Temporal Sulcus. Neuron, 41(5), 809–823. 10.1016/S0896-6273(04)00070-4

Beauchamp, M. S., Nath, A. R., & Pasalar, S. (2010). fMRI-Guided transcranial magnetic stimulation reveals that the superior temporal sulcus is a cortical locus of the McGurk effect. Journal of Neuroscience, 30(7), 2414–2417.

Benevento, L. A., Fallon, J., Davis, B. J., & Rezak, M. (1977). Auditory-visual interaction in single cells in the cortex of the superior temporal sulcus and the orbital frontal cortex of the macaque monkey. Experimental Neurology, 57(3), 849–872. 10.1016/0014-4886(77)90112-1

Bradski, G. (2000). The opencv library. Dr. Dobb’s Journal: Software Tools for the Professional Programmer, 25(11), 120–123.

Bruce, C., Desimone, R., & Gross, C. G. (1981). Visual properties of neurons in a polysensory area in superior temporal sulcus of the macaque. Journal of Neurophysiology, 46(2), 369–384. 10.1152/jn.1981.46.2.369

Calvert, G. A., Brammer, M. J., Bullmore, E. T., Campbell, R., Iversen, S. D., & David, A. S. (1999). Response amplification in sensory-specific cortices during crossmodal binding. Neuroreport, 10(12), 2619–2623. 10.1097/00001756-199908200-00033

Calvert, G. A., Campbell, R., & Brammer, M. J. (2000). Evidence from functional magnetic resonance imaging of crossmodal binding in the human heteromodal cortex. Current Biology, 10(11), 649–657.

Chen, J., Leong, Y. C., Honey, C. J., Yong, C. H., Norman, K. A., & Hasson, U. (2017). Shared memories reveal shared structure in neural activity across individuals. Nature Neuroscience, 20(1), 115–125. 10.1038/nn.4450

Devlin, J. T., & Poldrack, R. A. (2007). In praise of tedious anatomy. NeuroImage, 37(4), 1033–1041. 10.1016/j.neuroimage.2006.09.055

Diehl, M. M., & Romanski, L. M. (2014). Responses of prefrontal multisensory neurons to mismatching faces and vocalizations. The Journal of neuroscience : the official journal of the Society for Neuroscience, 34(34), 11233–11243. 10.1523/JNEUROSCI.5168-13.2014

Gavrilov, N., Hage, S. R., & Nieder, A. (2017). Functional Specialization of the Primate Frontal Lobe during Cognitive Control of Vocalizations. Cell Reports, 21(9), 2393–2406. 10.1016/j.celrep.2017.10.107

Gemba, H., Miki, N., & Sasaki, K. (1997). Cortical Field Potentials Preceding Vocalization in Monkeys. Acta Oto-Laryngologica, 117(sup532), 96–98. 10.3109/00016489709126151

Hasson, U., Nir, Y., Levy, I., Fuhrmann, G., & Malach, R. (2004). Intersubject Synchronization of Cortical Activity During Natural Vision. Science, 303(5664), 1634–1640. 10.1126/science.1089506

Huth, A. G., de Heer, W. A., Griffiths, T. L., Theunissen, F. E., & Gallant, J. L. (2016). Natural speech reveals the semantic maps that tile human cerebral cortex. Nature, 532(7600), 453–458. 10.1038/nature17637

Insausti, R. (2013). Comparative neuroanatomical parcellation of the human and nonhuman primate temporal pole. Journal of Comparative Neurology, 521(18), 4163–4176. 10.1002/cne.23431

Jääskeläinen, I. P., Sams, M., Glerean, E., & Ahveninen, J. (2021). Movies and narratives as naturalistic stimuli in neuroimaging. NeuroImage, 224. 10.1016/j.neuroimage.2020.117445

Khandhadia, A. P., Murphy, A. P., Romanski, L. M., Bizley, J. K., & Leopold, D. A. (2021). Audiovisual integration in macaque face patch neurons. Current Biology: CB, 31(9), 1826–1835.e3. 10.1016/j.cub.2021.01.102

McGurk, H., & MacDonald, J. (1976). Hearing lips and seeing voices. Nature, 264(5588), 746–748. 10.1038/264746a0

Morán, I., Perez-Orive, J., Melchor, J., Figueroa, T., & Lemus, L. (2021). Auditory decisions in the supplementary motor area. Progress in Neurobiology, 202, 102053. 10.1016/j.pneurobio.2021.102053

Niu, M., & Palomero-Gallagher, N. (2023). Architecture and connectivity of the human angular gyrus and of its homolog region in the macaque brain. Brain Structure & Function, 228(1), 47–61. 10.1007/s00429-022-02509-7

Peelle, J. E. (2014). Methodological challenges and solutions in auditory functional magnetic resonance imaging. Frontiers in Neuroscience, 8. 10.3389/fnins.2014.00253

Peelle, J. E., Spehar, B., Jones, M. S., McConkey, S., Myerson, J., Hale, S., Sommers, M. S., & Tye-Murray, N. (2022). Increased Connectivity among Sensory and Motor Regions during Visual and Audiovisual Speech Perception. The Journal of Neuroscience, 42(3), 435–442. 10.1523/JNEUROSCI.0114-21.2021

Perrodin, C., Kayser, C., Logothetis, N. K., & Petkov, C. I. (2014). Auditory and visual modulation of temporal lobe neurons in voice-sensitive and association cortices. The Journal of Neuroscience, 34(7), 2524–2537. 10.1523/jneurosci.2805-13.2014

Plakke, B., Hwang, J., & Romanski, L. M. (2015). Inactivation of Primate Prefrontal Cortex Impairs Auditory and Audiovisual Working Memory. The Journal of neuroscience : the official journal of the Society for Neuroscience, 35(26), 9666–9675. 10.1523/JNEUROSCI.1218-15.2015

Poremba, A., Saunders, R. C., Crane, A. M., Cook, M., Sokoloff, L., & Mishkin, M. (2003). Functional mapping of the primate auditory system. Science, 299(5606), 568–572. 10.1126/science.1078900

Romanski, L. M. (2007). Representation and integration of auditory and visual stimuli in the primate ventral lateral prefrontal cortex. Cerebral Cortex, 17 Suppl 1(Suppl 1), i61-69. 10.1093/cercor/bhm099

Romanski, L. M. (2012). Integration of faces and vocalizations in ventral prefrontal cortex: Implications for the evolution of audiovisual speech. Proceedings of the National Academy of Sciences of the United States of America, 109 Suppl 1(Suppl 1), 10717–10724. 10.1073/pnas.1204335109

Rozzi, S., Calzavara, R., Belmalih, A., Borra, E., Gregoriou, G. G., Matelli, M., & Luppino, G. (2006). Cortical Connections of the Inferior Parietal Cortical Convexity of the Macaque Monkey. Cerebral Cortex, 16(10), 1389–1417. 10.1093/cercor/bhj076

Saarimäki, H. (2021). Naturalistic Stimuli in Affective Neuroimaging: A Review. Frontiers in Human Neuroscience, 15, 675068. 10.3389/fnhum.2021.675068

Shamay-Tsoory, S. G., & Mendelsohn, A. (2019). Real-Life Neuroscience: An Ecological Approach to Brain and Behavior Research. Perspectives on Psychological Science, 14(5), 841–859. 10.1177/1745691619856350

Sherafati, A., Bajracharya, A., Jones, M. S., Speh, E., Munsi, M., Lin, C.-H. P., Fishell, A. K., Hershey, T., Eggebrecht, A. T., Culver, J. P., & Peelle, J. E. (2025). A high-density diffuse optical tomography dataset of naturalistic viewing. bioRxiv, 2023.11.07.565473. 10.1101/2023.11.07.565473

Smith, S. M., Jenkinson, M., Woolrich, M. W., Beckmann, C. F., Behrens, T. E., Johansen-Berg, H., … & Matthews, P. M. (2004). Advances in functional and structural MR image analysis and implementation as FSL. Neuroimage, 23, S208–S219.

Smith, S. M., & Nichols, T. E. (2009). Threshold-free cluster enhancement: addressing problems of smoothing, threshold dependence and localisation in cluster inference. Neuroimage, 44(1), 83–98.

Sonkusare, S., Breakspear, M., & Guo, C. (2019). Naturalistic Stimuli in Neuroscience: Critically Acclaimed. Trends in Cognitive Sciences, 23(8), 699–714. 10.1016/j.tics.2019.05.004

Stevenson, R. A., VanDerKlok, R. M., Pisoni, D. B., & James, T. W. (2011). Discrete neural substrates underlie complementary audiovisual speech integration processes. NeuroImage, 55(3), 1339–1345. 10.1016/j.neuroimage.2010.12.063

Sumby, W. H., & Pollack, I. (1954). Visual Contribution to Speech Intelligibility in Noise. The Journal of the Acoustical Society of America, 26(2), 212–215. 10.1121/1.1907309

Van Engen, K. J., Dey, A., Sommers, M. S., & Peelle, J. E. (2022). Audiovisual speech perception: Moving beyond McGurk. The journal of the acoustical society of america, 152(6), 3216–3225.

Von Lühmann, A., Zheng, Y., Ortega-Martinez, A., Kiran, S., Somers, D. C., Cronin-Golomb, A., Awad, L. N., Ellis, T. D., Boas, D. A., & Yücel, M. A. (2021). Toward Neuroscience of the Everyday World (NEW) using functional near-infrared spectroscopy. Current Opinion in Biomedical Engineering, 18, 100272. 10.1016/j.cobme.2021.100272

